# Acceleration of genome rearrangement in clitellate annelids

**DOI:** 10.1101/2024.05.12.593736

**Authors:** Darrin T. Schultz, Elizabeth A.C. Heath-Heckman, Christopher J. Winchell, Dian-Han Kuo, Yun-sang Yu, Fabian Oberauer, Kevin M. Kocot, Sung-Jin Cho, Oleg Simakov, David A. Weisblat

## Abstract

Comparisons of multiple metazoan genomes have revealed the existence of ancestral linkage groups (ALGs), genomic scaffolds sharing sets of orthologous genes that have been inherited from ancestral animals for hundreds of millions of years (Simakov et al. 2022; Schultz et al. 2023) These ALGs have persisted across major animal taxa including Cnidaria, Deuterostomia, Ecdysozoa and Spiralia. Notwithstanding this general trend of chromosome-scale conservation, ALGs have been obliterated by extensive genome rearrangements in certain groups, most notably including Clitellata (oligochaetes and leeches), a group of easily overlooked invertebrates that is of tremendous ecological, agricultural and economic importance (Charles 2019; Barrett 2016). To further investigate these rearrangements, we have undertaken a comparison of 12 clitellate genomes (including four newly sequenced species) and 11 outgroup representatives. We show that these rearrangements began at the base of the Clitellata (rather than progressing gradually throughout polychaete annelids), that the inter-chromosomal rearrangements continue in several clitellate lineages and that these events have substantially shaped the evolution of the otherwise highly conserved Hox cluster.

## INTRODUCTION

While changes in developmental processes are the immediate/proximal cause of changes in body plan during evolution, the underlying/ultimate cause is changes in the genomic information that is largely responsible for programming the developmental processes. A broad spectrum of genomic changes is possible, ranging from relatively frequent (point mutations and inversions) to much rarer (gene duplications, chromosomal translocations and whole genome duplications). Large differences in the frequency of successful (inherited) inversions versus translocations result in the observation that, over large evolutionary timescales, gene co-linearity among species is lost while chromosomal synteny is preserved, i.e. orthologous genes among species move back and forth along their respective chromosomes while largely maintaining their chromosomal identity, even among species as divergent as cnidarians and bilaterians, that have been evolving independently for over 600 MY (Simakov et al. 2022). Thus, conservation of synteny has allowed the inference of 29 ancestral chromosomes/chromosome arms (often referred to as ancestral linkage groups, ALGs) in the last common metazoan ancestor (Simakov et al. 2022; Schultz et al. 2023).

Exceptions to syntenic conservation exist, however. In particular, when the first three spiralian genomes were sequenced (Simakov et al. 2013), the mollusc *Lottia* and the polychaete annelid *Capitella* [separated by 534-636 MY (dos Reis et al. 2015)] showed strong synteny, but the genome of the leech *Helobdella* [a clitellate annelid separated by 476-636 MY from *Capitella* (dos Reis et al. 2015) showed no synteny with either species; the dramatic acceleration of genome rearrangements in *Helobdella* extended even to the highly conserved Hox cluster, which exhibits multiple duplications and deletions along with atomization of the ancestral spiralian 11 gene cluster itself relative to *Lottia* and *Capitella*.

This dramatic loss of chromosomal synteny is of interest for several reasons. A priori, the relaxation of evolutionary constraints on genome architecture enhances evolutionary rates and may contribute to the evolution of novelty at genomic and morphological levels. The mechanisms by which synteny is lost are also of considerable interest. Inter-chromosomal translocations are frequently lethal because they can lead to an aberrant pairing of chromosomes during meiosis and/or gene dosage defects in the resulting gametes and embryos (Wright 1941). How might this problem have been circumvented in the lineage leading to *Helobdella*? Do the accelerated genome rearrangements result from an acceleration in all aspects of genome evolution for this taxon, or a specific relaxation of constraints on genome organization? Do the rearrangements seen in *Helobdella* result from a transient relaxation of constraints on genome organization followed by fixation somewhere in the lineage leading to *Helobdella*, perhaps within the Clitellata, or do they reflect an ongoing process?

To address these questions, we have compared the organization of 23 spiralian taxa for which well-assembled genomes are currently available (Figure 1A): 12 clitellate annelids, including 4 newly sequenced species (Figure 1B); 9 polychaete annelids; and 2 molluscan outgroups. We confirm a dramatic loss of chromosomal synteny between all the clitellate species with respect to polychaete and molluscan species. Even the most closely related polychaete species (*Capitella teleta*) sampled here shows no increased loss of ALGs (Simakov et al. 2022). Moreover, this loss of synteny reflects an acceleration of genome rearrangements and is accompanied by elevated rates of protein sequence evolution among several of these taxa. Finally, comparisons among the available clitellate genomes suggest that the acceleration of genome rearrangements reflects an ongoing relaxation of constraints on genome organization, rather than a transient event at the base of the Clitellata. We speculate that this change results from a combination of developmental, physiological and/or ecological factors associated with the invasion of freshwater and terrestrial habitats by clitellate annelids.

**Figure 1:**
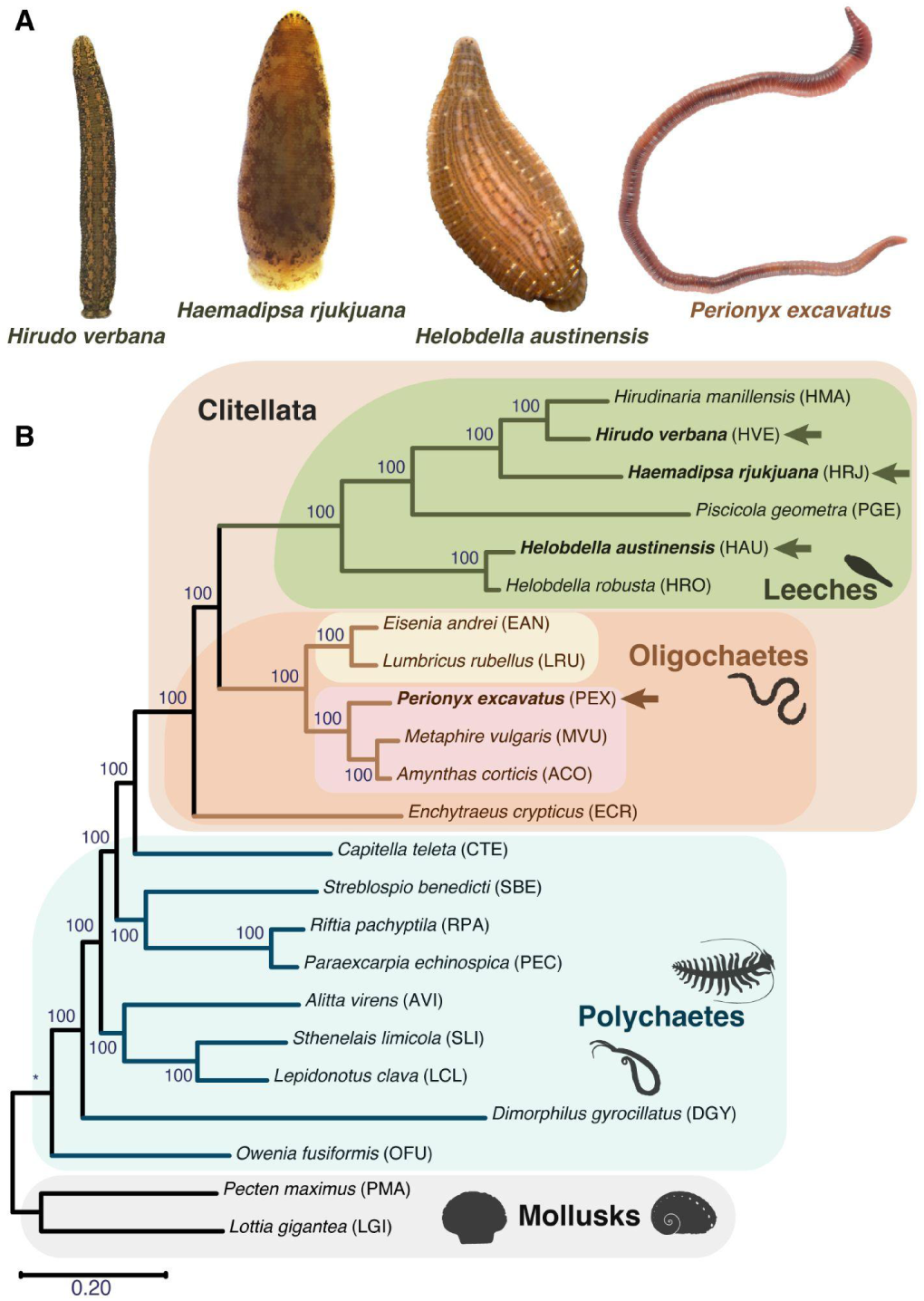
New clitellate genomes and clitellate-annelid relationships. **A.** Four newly sequenced cliitellate annelids (dorsal views, anterior is up). *Hirudo verbana* (HVE, small adult 4.5, cm long). *Haemadipsa rjukjuana* (HRJ, adult, 2 cm long). *Helobdella austinensis* (HAU, adult, 1.5 cm long). *Perionyx excavatus* (PEX, adult, 11 cm long). **B.** Phylogenetic relationships (Maximum Likelihood tree based on 250 orthologous genes) among the 23 species (2 mollusks; 9 polychaete annelids; and 12 clitellate annelids-6 oligochaetes and 6 leeches) used in this study (see Materials and Methods for details). Each species is assigned a three letter identifier. Photography credits: *H. verbena* and *H. austinensis* by Christopher J. Winchell © 2024. *H. rjukjuana* and *P. excavatus* by Sung-Jin Cho © 2024.

## RESULTS AND DISCUSSION

### Four new annelid genomes

*Helobdella austinensis* (HAU) is a small glossiphoniid leech, first identified in Austin Texas, USA (Kutschera et al. 2013), that feeds on freshwater snails and other aquatic invertebrates. It is a sibling species to *H. robusta* (HRO), which was among the first three spiralians to be sequenced (Simakov et al. 2013). HAU was sequenced because it largely replaced HRO as an experimental model due to its resistance to die-offs in lab culture. 90% of the 187 Mb Mb genome was assembled onto 41 scaffolds ≥1 Mb in length, using Illumina sequencing and HiRise assembly pipeline (Dovetail).

*Haemadipsa rjukjuana* (HRJ) is a sanguivorous land leech found in Taiwan, Japan and Korea (Lai, Nakano, and Chen 2011; Won et al. 2014). To our knowledge, this is the first haemadipsid leech species to be sequenced. Chromosome-level genome assembly analysis was achieved using PacBio sequencing and proximity ligation (Hi-C, Dovetail). The 139.8 Mb genome was assembled into 33 scaffolds, with 99% of the assembly on 11 chromosomes.

*Hirudo verbana* (HVE) is a hirudinid leech, one of two species of European medicinal leech (Siddall et al. 2007); it is widely used for the analysis of neural circuits underlying behavior, and by plastic surgeons for relief of vasocongestion resulting from reconstructive surgeries (Siddall et al. 2007; Kraemer et al. 1988; Houschyar et al. 2015). Previous analyses of its genome assemblies (Babenko et al. 2020; Kvist et al. 2020) had yielded short scaffolds. Here, a combination of PacBio sequencing and proximity ligation (Dovetail) yielded near chromosomal scale assembly. Roughly half of the 194 Mb genome was assembled onto 7 scaffolds, and 90% was assembled onto 35 scaffolds ≥1 Kb in length.

*Perionyx excavatus* (PEX) is a megascolecid earthworm. It probably originated from the Indian subcontinent, but it is commonly used in vermicomposting and has been collected at locations around the world (Hendrix et al. 2008). A chromosome-level genome assembly was achieved using PacBio sequencing and proximity ligation (Hi-C, Dovetail). The genome was assembled into 645 scaffolds with a size of 837 Mb, and 22 scaffolds corresponding to chromosomes were larger than 10 Mb, with a coverage of 92.16%.

### Widespread loss of chromosomal synteny among clitellate annelids

Our analysis confirms the previous notion that chromosomes are generally conserved in the animal kingdom and can be represented by “algebraic” combinations of 29 ancestral animal linkage groups (Hendrix et al. 2008; Simakov et al. 2022). While the overall fusion rates can be different among animal clades, the overall background rate is relatively low among most of the metazoans (few fusions per hundreds of millions of years). However, some lineages show ‘peaks’ in such rearrangements that seem to occur in a time-restricted manner. For example, at the base of coleoid cephalopods, there has been a large-scale ALG reshuffling that resulted in a new set of linkage groups that have remained comparatively stable so far and only underwent species-specific fusions (Albertin et al. 2022).

The well-conserved synteny that is evident in mollusks and polychaete annelids has also been lost for all the clitellate genomes surveyed here (Figure 2). Our genomic sampling suggests that the pace of genome rearrangements was dramatically accelerated at some point in the lineage leading to earthworms and leeches after it diverged from the capitellid lineage. In contrast to most other taxa, however, our findings for clitellates are striking in that not only the ancestral metazoan linkage group complement has been scrambled, but the inter-chromosomal scrambling may still be ongoing in at least some of the clitellate lineages (Figure 3).

Additional translocation-rich events happened at the base of each of the major clitellate lineages, including Megascolecidae, Lumbricidae, and Hirudinida. Rearrangements are also evident within the recent branches of these taxa. For example, in the leech lineages, multiple species-specific translocations can be observed. This enhanced rate of rearrangements is accompanied by an increased rate of protein sequence evolution among leeches (Figure 1A). Our findings are thus more consistent with a maintained acceleration of rearrangements in this group rather than a transient burst of accelerated rearrangements at the base followed by a return to relative stasis.

On top of the enhanced inter-chromosomal translocation rate, our analysis also detected whole genome duplications, confirming the already reported duplication in *Metaphire* (Jin et al. 2020). We observed the same duplicated karyotype in *Amynthas*, but not in *Perionyx*, which narrows the likely timing of this duplication event. Together, these data put clitellates forward as a model system to study the effect of chromosomal rearrangements in evolution.

**Figure 2:**
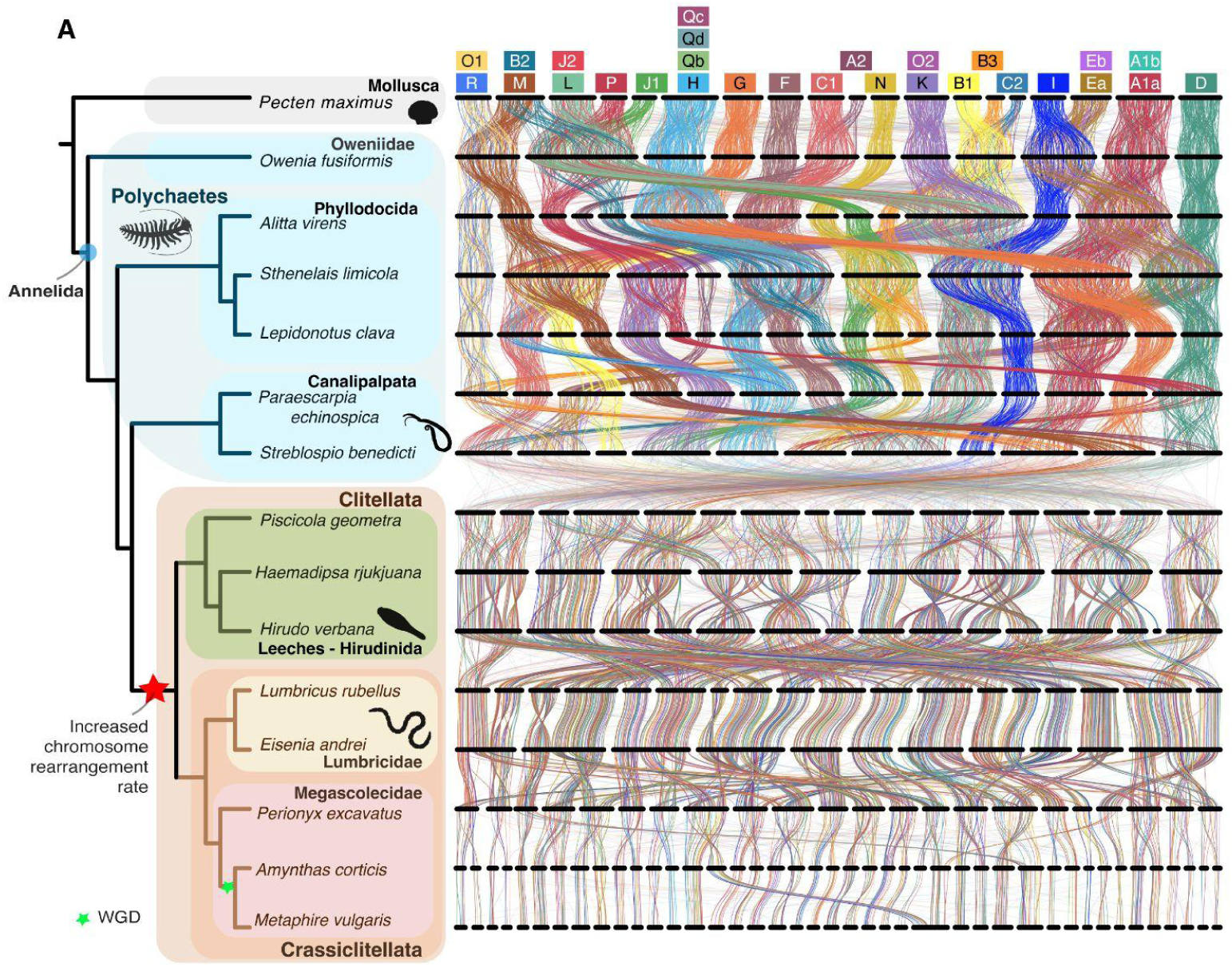
Rapid genome rearrangement in clitellate annelids. **A.** Synteny plot showing orthologs (curved vertical lines) on chromosomal scaffolds (horizontal black bars) between pairs of species. Orthologs are colored based on bilaterian-cnidarian-sponge ancestral linkage groups (BCnS ALGs). Orthologs on significantly-related chromosomes are opaque (Fisher’s exact test ≤ 0.05), and orthologs on non-significant chromosome pairs are translucent. All major annelid clades, except the Clitellata, retain the BCnS ALGs that are conserved in other metazoans, including the scallop *Pecten maximus*. The genomes of extant clitellates show that there have been extensive structural rearrangements in the branch leading to clitellates, and within the clitellates.

**Figure 3:**
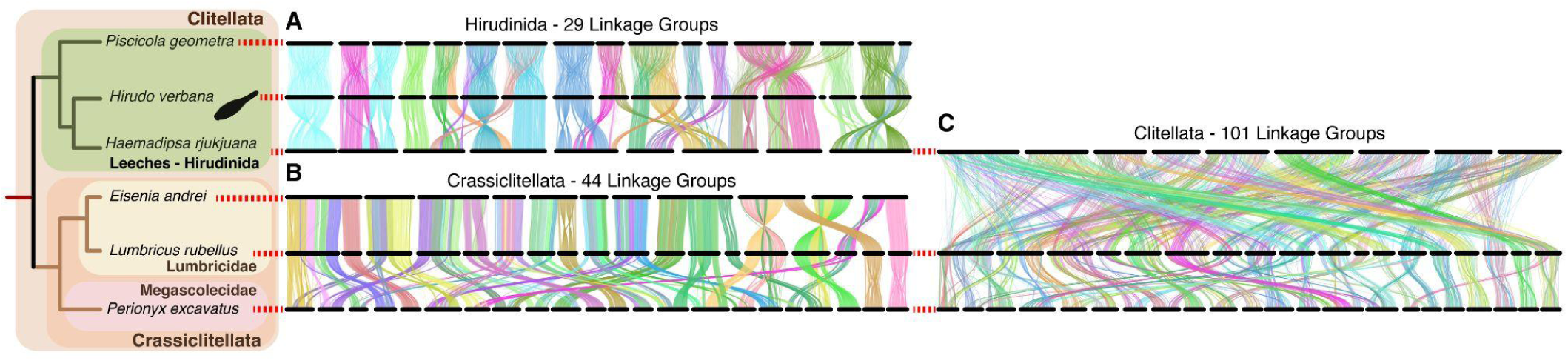
Increased inter-chromosomal rearrangements in clitellate annelids. All three panels show significantly large linkage groups from species trios, inferred with the program `odp_nway_rbh`. The colors do not correspond between the panels. **A.** Ribbon diagram colored by the 29 linkage groups inferred from the clade Hirudinida. **B.** Ribbon diagram colored by the 44 linkage groups inferred from the clade Crassiclitellata. **C.** The ribbon diagram showing members of both the Hirudinida, the Lumbricidae, and the Megascolecidae. The synteny plot showing orthologs (curved vertical lines) on chromosomal scaffolds (horizontal black bars). There are not only rearrangements between the deep Hirudinida-Crassiclitellata node, but among more recently diverged species.

### Atomization of the Hox cluster among clitellates suggests that clitellate genome rearrangements are ongoing

Previous work has shown that Hox genes in *Helobdella robusta* are not organized in a clustered and linear array on a chromosome, as observed in many bilaterian genomes. Instead, they are broken into a few subclusters and singlets, with a number of duplicates and gene losses (Simakov et al. 2013). This “atomized” Hox configuration was observed in all clitellate species examined here; furthermore, they differ in various degrees among species (Supplementary Table 1).

To determine the extent of Hox gene translocations in the extensively rearranged clitellate genomes, we compared Hox inventories on syntenic scaffolds in three separate clitellate subgroups: lumbricids, megascolecids, and leeches. For each subgroup, we chose one anchor species to which other subgroup members were compared. The anchor species was chosen as the one with the fewest Hox-containing genome scaffolds (*Eisenia* for lumbricids and *Haemadipsa* for leeches), or as the taxon presumed to branch basally among sampled members of a subgroup (*Perionyx* for megascolecids). We grouped each Hox-containing scaffold in the anchor species with syntenic scaffolds of other subgroup members into “synteny sets” (Supplementary Table 1).

Comparing the Hox inventories within synteny sets allowed us to differentiate whether syntenic clitellate chromosomes contain conserved sub-clusters or random assortments of Hox genes (Figure 4). For the lumbricid oligochaetes (represented by *Lumbricus* and *Eisenia*), three out of six synteny sets have identical Hox inventories (Supplementary Table 1): set 4 (Hox3, two Scr, Post2), set 5 (Lab, Hox3, Scr, Lox5), and set 6 (Post1). In synteny set 1, each genus possesses Pb and Post2, but they differ with respect to a third gene, which in *Eisenia* is Scr and in *Lumbricus* is an additional copy of Post2. Both genera possess 12 genes in synteny set 3, but relative to *Eisenia*, *Lumbricus* is missing three copies of Lox4 and two copies of Scr, has gained an additional copy of Lab and of Post2, and possesses two copies of Dfd and one of Lox2 (Dfd and Lox2 were not found on any *Eisenia* scaffold).

Within megascolecid oligochaetes, *Metaphire* and *Amynthas* are within the Pheretima complex, a morphologically derived group within Megascolecidae (“Origin and Diversification of Pheretimoid Megascolecid Earthworms in the Japanese Archipelago as Revealed by Mitogenomic Phylogenetics” 2023; Sims and Easton 1972) Sato et al. 2023(“Origin and Diversification of Pheretimoid Megascolecid Earthworms in the Japanese Archipelago as Revealed by Mitogenomic Phylogenetics” 2023; Sims and Easton 1972), and exhibit a shared whole genome duplication relative to *Perionyx* (Figure 2). Thus: 1), most individual chromosomes in *Perionyx* correspond to two paralogous chromosomes in *Amynthas* and in *Metaphire*, and 2) each of these paralogous chromosomes in *Amynthas* has an orthologous chromosome in *Metaphire.* Among these three megascolecid genomes, the Hox cluster remnants on the paralogous and orthologous chromosomes exhibit similarities consistent with their respective ancestries, but also differences, suggesting recent or ongoing inter-chromosomal translocations (Supplementary Table 1).

Thus, consistent with the whole genome duplication, we found that each of the eleven Hox-containing scaffolds in *Perionyx* is syntenic with two scaffolds in both *Metaphire* and *Amynthas* (Supplementary Table 1). The Hox inventories across 12 megascolecid synteny sets (Supplementary Table 1) were somewhat labile, however. In multiple cases, the presence of a particular Hox gene or set of Hox genes within a given synteny set was taxon-specific: *Perionyx* (Scr in set 1; Hox3 and Scr in set 3; Post2 in set 9; Post2, Lox2, and Lox4 in set 11), *Metaphire* (Pb in set 3; Pb, Lox2, Hox3 in set 6; Hox3 in set 12), *Amynthas* (Lox5 in set 6; Lox4 in set 7), and *Perionyx* plus *Metaphire* (Post2 in set 2; Scr in set 4; Lox2 in set 7; Scr and Hox3 in set 10). Despite these differences, we found that all three megascolecid species had in common the presence of: Pb and Post2 in set 1; Lab and Lox5 in set 3; Antp in set 4; Post1 in set 5; Post2, Lox4, Dfd, and Lab in set 6; Hox3 and Scr in set 7.

Finally, we used *Haemadipsa rjukjuana* as the anchor species for comparing Hox organization in leeches (represented by *Helobdella*, *Piscicola*, *Haemadipsa*, *Hirudo*) and found conservation of Hox inventory within all three synteny sets (Supplementary Table 1). Set 1, contains seven Hox genes in two apparent sub-clusters (Lab-Scr-Lox5 and Lab-Dfd-Lox4-Post2) in all four genera. *Piscicola* Hox3 is the only other Hox gene in leech synteny set one. Set 2, containing Hox3 and an apparent Lox2-Scr subcluster, is conserved in *Haemadipsa*, *Helobdella*, and *Hirudo* but only partially conserved in *Piscicola*, for which no Lox2 gene was found and, as mentioned previously, seems to have moved its Hox3 gene from set 2 to set 1. Set 3, containing up to 8 Hox genes, including an apparent Dfd-Lox4-Lox2-Post2 subcluster, Antp, and three Scr genes, is perfectly conserved in *Haemadipsa* and *Helobdella*, but *Piscicola* and *Hirudo* are each missing two Scr genes. Moreover, the set 3 inventory of *Hirudo* is unique in that it appears to have lost Lox2 and Antp. Consistent with previous reports, we found no evidence of Pb or Post1 in any of these leech genomes (Simakov et al. 2013).

**Figure 4:**
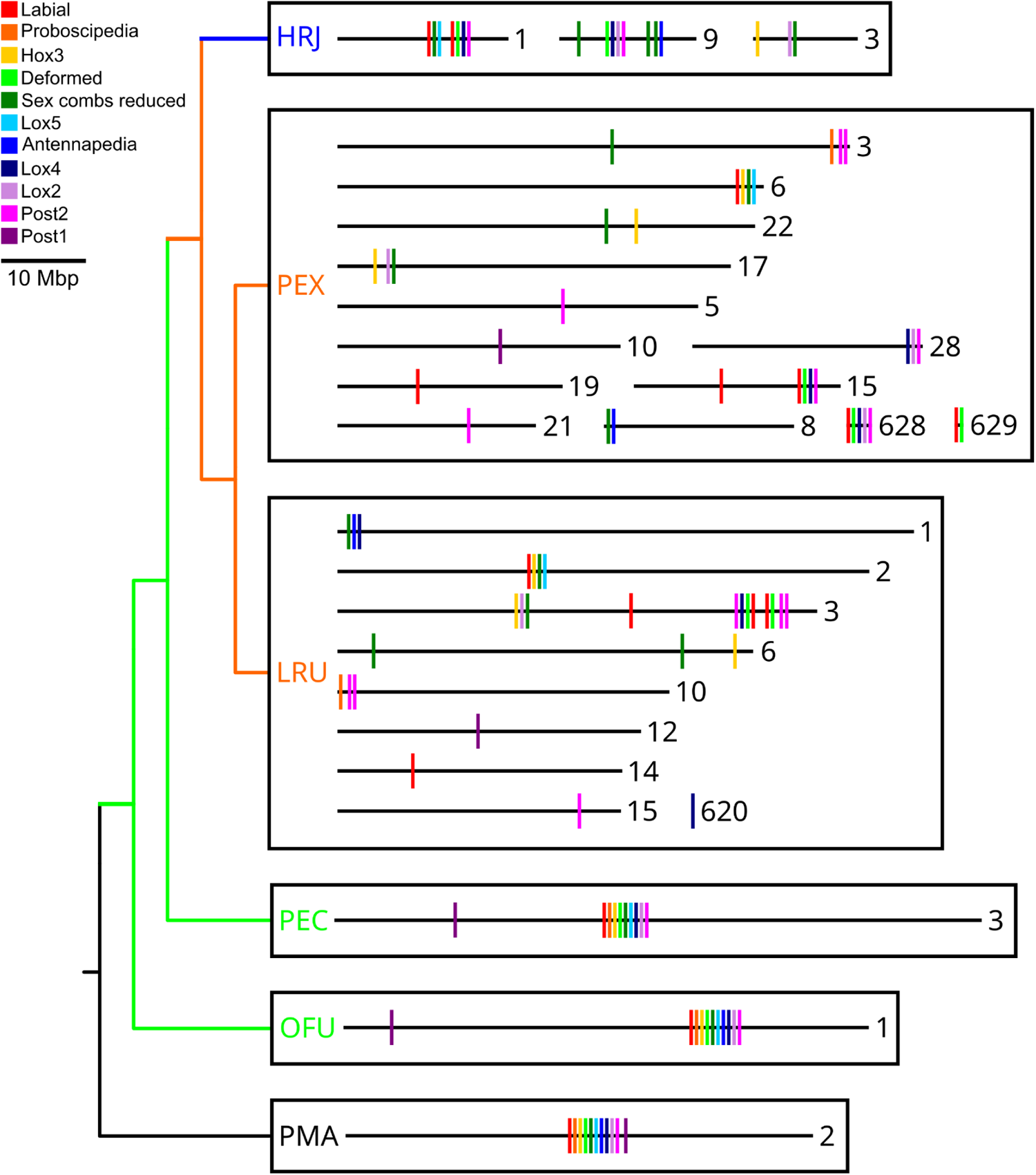
Dispersal of the Hox Cluster, exemplified in selected oligochaetes (orange) and leeches (blue), relative to polychaete (green) and molluscan (black) outgroups. Numbers refer to Hox-containing genome scaffolds, which are drawn to scale. Hox genes (colored vertical lines on scaffolds) are not to scale. Taxon names: PMA, *Pecten maximus*; OFU, *Owenia fusiformis*; PEC, *Paraescarpia echinospica*; LRU, *Lumbricus rubellus*; PEX, *Perionyx excavatus*; HRJ, *Haemadipsa rjukjuana*.

### Potentially conserved Hox subclusters between oligochaetes and leeches

We also observed distinct associations of several sets of Hox genes across clitellate subgroups defined by their consistent colocalization on the same chromosomes.

1. Lab, Dfd, Lox4, and Post2 are contiguous within **lumbricid** synteny set 3 (*Lumbricus* only, as *Eisenia* is missing Dfd), **megascolecid** synteny set 6, and **leech** synteny set 1. The chromosomes in these sets share some, but not much, synteny between subgroups.
2. Hox3, Scr, and Lox2 are contiguous within **megascolecid** synteny set 7, **leech** synteny set 2, and possibly **lumbricid** synteny set 3.
3. Lab, Scr, and Lox5 (plus Hox3 in oligochaetes) are contiguous within **lumbricid** synteny set 5, **megascolecid** synteny set 3 (*Perionyx* only), and **leech** synteny set 1.
4. Scr and Antp are contiguous within **megascolecid** synteny set 4, **leech** synteny set 3, and possibly **lumbricid** synteny set 1 (Scr and Antp are unplaced in *Eisenia* but occur with Lox4 on *Lumbricus* scaffold 1).

### Hox gene duplications and losses

As detailed above, the atomization of the Hox complement in clitellate annelids has been accompanied by numerous instances of gene duplication. In leeches, we also note two examples of Hox gene loss, namely *pb*, an anterior group gene (second in the ancestral cluster), and *post1* (a posterior group gene that comes last in the ancestral cluster).

A hallmark in the divergence of oligochaetes (earthworms) and true leeches from polychaetes is the reduction and then total loss of segmentally iterated bristles (chaetae) that serve to reduce slippage during peristaltic locomotion that is so familiar from earthworms. Within the canonical spiralian Hox cluster, *post1* occupies the terminal position of the posterior group and is expressed in the chaetal sacs of several annelid and brachiopod species (Fröbius, Matus, and Seaver 2008; Kulakova et al. 2007; Schiemann et al. 2017) and is thought to play an integral role in chaetae function and development. Interestingly, we note that Post1 is somewhat disproportionately separated from the rest of the Hox cluster in the bivalve mollusc (*Pecten maximus* - PMA, Figure 4)--specifically, the ratio of the length of the full 11 gene cluster to the length of the 10 gene cluster when *post1* is excluded is 1.4. In *Owenia fusiformis* - OFU, representing the deepest branch of polychaete annelids in this study, this ratio is increased to 78.2 and *post1* is now located upstream of the anterior group genes, indicative of either an inversion or transposition. The isolation of *post1* from other Hox genes is further increased in the clitellates–*post1* is commonly the only Hox gene on its scaffold in oligochaetes and is missing altogether from leech chromosomes. The absence of *post1* in the leeches is coincidental with their loss of chaetae. Further investigation into the role and consequence of *post1* and associated genes in chaetae-bearing annelids may help clarify whether clitellates lost chaetae due to a genomic deletion of *post1*, or if *post1* was lost by another function, such as relaxation of selective constraint after the loss of function at other loci. A second evolutionary loss from the leech species examined here is the anterior group Hox gene *pb*, for which there is no obvious morphological correlate.

### Factors that may contribute to relaxation of constraints on genome rearrangements in Clitellata

It is clear from our synteny analyses that the rate of genome rearrangements (more precisely, the inheritance and persistence of rearrangements that we measure in extant species) was elevated in the evolutionary branches leading up to, and within, the Clitellata. The protein phylogenomic results show that, despite the increased rate of genomic structural changes, the rate of protein evolution in the clitellate crown group is not particularly elevated relative to other annelids. The inference that the rate of genome rearrangements *remained* elevated within this group after initial divergence from all other annelids suggests that it does not reflect a transient process such as a wave of transposon activity that has since been silenced. The analysis of repetitive elements in the *Helobdella* genome also argues against that explanation (Simakov et al. 2013) and no effect on syntenic preservation in extremely repeat-rich genomes in vertebrates have been observed, e.g., in the lungfish (Meyer et al. 2021). The population-level or chromosome-scale sequencing of more genomes within the clitellates, and more broadly the annelids, will clarify the contemporary genome rearrangement rates in these clades.

We speculate that environmental factors associated with the emergence of clitellate annelids may have enabled the acceleration of heritable genome rearrangements in this clade. Freshwater and terrestrial habitats occupied by clitellates are apt to be less stable than the marine environment in which their polychaete ancestors evolved (Kuo, 2017). More ephemeral habitats may be expected to result in smaller and less stable populations, and smaller effective population size in turn facilitates the fixation of genomic changes through genetic drift (Viard, Justy, and Jarne 1997; Charbonnel et al. 2002). Another possible environmental factor contributing to increased rates of genome arrangement would be the presence of locally concentrating clastogenic agents (e.g., irradiation or chemicals) in the clitellate habitats, which are more heterogeneous and fragmented than the ancestral marine habitats.

We also speculate that evolutionary innovations associated with the emergence of clitellate annelids, namely simultaneous hermaphroditism, and self-fertilization in some taxa, might have concomitantly increased the rate at which genome rearrangements became fixed. Simultaneous hermaphroditism occurs in clitellate annelids, but is rare in polychaetes. The advantage of being able to mate with any con-specific adult in a small and/or low density population presumably offsets the energetic/physiological cost of developing two sets of reproductive organs (Heath, 1977). A more extreme adaptation to infrequent mating encounters is the capacity for self-fertilization. Self-mating, and to a lesser extent sibling matings, provide a possibility of rescuing chromosomal translocations that would otherwise prove lethal through defects in homolog pairing and/or gene dosage effects.

Finally, we previously identified developmental considerations that might also contribute to the observed increase in genome rearrangements. In many animal species, dedicated germline precursor cells (PGCs) arise early in the embryonic cell divisions, which reduces the number of mitoses required to get from the zygote to the gametes that will found the next generation. For example, PGCs in at least some polychaetes arise as the first two cells produced by the bilateral pairs of mesodermal stem cells, corresponding to eight and nine rounds of zygotic mitoses from the fertilized egg in the conserved spiralian cleavage patterns [reviewed by (Rebscher 2014)]. In contrast, the PGCs in clitellate embryos are not segregated from somatic lineages until after at least 24 rounds of zygotic mitoses and male and female PGC precursors share the same lineages through at least 23 rounds (Rebscher 2014; Kang, Pilon, and Weisblat 2002); (Oyama and Shimizu 2007; Cho, Vallès, and Weisblat 2014)--this long cell lineage delay to the segregation of PGCs increases both the opportunities for chromosomal aberrations to occur and the likelihood that any chromosomal alterations will be shared by male and female PGCs. Thus, we suggest that the increased rate of genome rearrangements that we have documented here for clitellate annelids could result from an increased probability for generating chromosomal abnormalities that are shared between the male and female gametes, combined with the capacity for rescuing such abnormalities through self-fertilization or sibling mating.

We note that other simultaneously hermaphroditic species are known, including mangrove killifish, and pulmonate molluscs, but sequenced representatives of these taxa do not appear to exhibit accelerated genome rearrangements (Supplementary Figure 1). Comparisons among these diverse taxa with respect to such variables population size and developmental details of PGC formation may provide further insights into the mechanisms of genome rearrangement acceleration.

## CONCLUSIONS

In this manuscript, we report a comparison of key spiralian and in particular clitellate taxa in terms of their phylogeny and chromosomal evolution. Our analyses confirm the faster rates of protein sequence evolution in the leech lineages. Chromosomal-level synteny comparisons reveal that the clitellate lineage underwent substantial chromosomal reorganizations, fueled by almost complete rearrangement of the ancestral metazoan linkage groups. While such genome-wide events were likely time-restricted, some lineages (e.g., leeches) kept up a higher translocation rate between chromosomes after their initial divergence from other annelids. This is also reflected in the highly partitioned and partially duplicated Hox clusters. While the mechanisms behind the breakage of ancestral chromosomal synteny in animal genomes remain elusive, this paper identifies several key biological factors that may have driven such events in clitellates, including self-mating and late germline separation.

## Supporting information

Supplemental Table 1 - docx format

Supplemental Table 1 - pdf format

## ACKNOWLEDGEMENTS

SJC acknowledges support by the Basic Science Research Program through the National Research Foundation of Korea (NRF) funded by the Ministry of Education (2020R1A6A1A06046235, 2023R1A2C1007868), and by grants from the National Institute of Biological Resources (NIBR), funded by the Ministry of Environment (MOE) of the Republic of Korea (NIBR202305101, NIBR202333201). DAW and DHK acknowledge support from Human Frontiers Science Program Grant RGP0060/2019. DTS and OS were supported by the European Research Council’s Horizon 2020: European Union Research and Innovation Programme, grant no. 945026. EACH was supported by the National Institute of General Medical Sciences of the NIH under Award Number R35GM150478. The computational results of this work have been supported by using the Life Science Compute Cluster (LiSC) of the University of Vienna.

## METHODS

### Genome database preparation

The following chromosome-scale and scaffold-level annelid and mollusk genomes, as well as their genome annotations and protein sequences, were downloaded for this project: *Helobdella robusta* (GCF_000326865.2) (Simakov et al. 2013), *Hirudinaria manillensis* (GCA_034509925.1) (Guan et al. 2019), *Metaphire vulgaris* (GCA_018105865.1) (Jin et al. 2020), *Amynthas corticis* (GCA_900184025.1) (Wang et al. 2021), *Eisenia andrei* (GWHACBE00000000) (Shao et al. 2020), *Enchytraeus crypticus* (GCA_905160935.1) (Amorim et al. 2021), *Capitella teleta* (GCA_000328365.1) (Simakov et al. 2013), *Streblospio benedicti* (GCA_019095985.1) (Zakas et al. 2022), *Dimorphilus gyrociliatus* (GCA_904063045.1) (Martín-Durán et al. 2021), *Owenia fusiformis* (GCA_903813345.2) (Martín-Zamora et al. 2023), *Lottia gigantea* (GCF_000327385.1) (Simakov et al. 2013), *Pecten maximus* (GCA_902652985.1) (Kenny et al. 2020), *Riftia pachyptila* (https://phaidra.univie.ac.at/detail/o:1220865) (de Oliveira et al. 2022), *Lepidonotus clava* (GCA_936440205.1) (Darbyshire et al. 2022), *Sthenelais limicola* (GCA_942159475.1) (Darbyshire et al. 2023), *Alitta virens* (GCA_932294295.1) (Fletcher et al. 2023), *Paraescarpia echinospica* (GCA_020002185.1) (Sun et al. 2021), *Lumbricus rubellus* (GCA_945859605.1) (Short et al. 2023), and *Piscicola geometra* (GCA_943735955.1) (Doe 2023). Additionally, we downloaded the following genomes of the simultaneously hermaphroditic species from NCBI: *Biomphalaria glabrata* (GCA_947242115.1) (The Darwin Tree of Life Project et al. 2022), and *Kryptolebias marmoratus* (GCF_001649575.2).

The genomes were prepared for use with the odp software package (Schultz et al. 2023) by extracting the protein coordinate information to rbh files.

### Sample collection

Adult earthworm *Perionyx excavatus* were obtained from the Nan-ji Water Reclamation Center (37° 35’ 13“ N, 126° 50’ 45” E) in Kyunggi-do, South Korea. Adult leech *Haemadipsa rjukjuana* were collected on Mt. Dock-Sil (altitude 639m) on the island Gageo-do (E 125° 07’, N 34° 04’), South Korea. The earthworms and leeches were collected in compliance with local regulations, and collected animals were maintained in an animal incubation room under controlled conditions. *Helobdella austinensis* (Kutschera et al. 2013) adults were originally collected from Shoal Creek in Austin, TX (within 1 kilometer of 30° 17’ 41“ N, 97° 44’ 44” W) in 1998 and have been in continuous laboratory culture since. The collection of *H. austinensis* leeches was performed in compliance with Texas regulations (TAC Title 31, PT 2 TPWD, Ch 69, Subchapter J, Rule §69.302). Adult *Hirudo* were obtained from a commercial supplier (Leeches.com).

### Haemadipsa rjukjuana and Perionyx excavatus methods

#### *H. rjukjuana* and *P. excavatus* DNA and RNA extraction

The MagAttract HMW DNA Kit (Qiagen, catalog no.67563) was used to extract DNA from the the muscle tissue was sampled from an adult *H. rjukjuana* with a body length of 2 cm, and an adult *P. excavatus* with a body length of 11 cm. DNA was eluted with 100ul of Solution AE. Total RNA was extracted from six individuals of each species using mirVana miRNA Isolation Kit (Ambion) according to the manufacturer’s recommended procedures.

#### *H. rjukjuana* and *P. excavatus* DNA Short-read Sequencing

We first used a NanoDrop spectrometer to confirm that the *H. rjukjuana* and *P. excavatus* DNA was pure, with an OD260/280 ratio of 1.8–2.0. We also verified that the DNA was intact, using a 1% agarose gel electrophoresis. Lastly, we quantified the DNA with a Qubit dsDNA HS Assay Kit (Thermo Fisher Scientific).

Whole-genome shotgun sequencing libraries were prepared according to the Illumina Truseq Nano DNA Library prep protocol. Briefly, 0.2µg of high molecular weight (HMW) DNA was randomly sheared using the Covaris S2 system to a median insert size of 350 basepairs. The fragmented DNA was prepared according to the manufacturer’s protocol, and the quality of the amplified libraries was verified by capillary electrophoresis (Bioanalyzer, Agilent). The libraries were quantified using qPCR with SYBR Green PCR Master Mix (Applied Biosystems), then were pooled in equimolar amounts. The pool was sequenced on an Illumina NovaSeq 6000 system following the standard Illumina protocols for 2x150 bp sequencing.

#### *H. rjukjuana* and *P. excavatus* RNA Short-read Sequencing

The *H. rjukjuana* and *P. excavatus* RNA purity was determined by assaying 1 µl of total RNA extract on a NanoDrop8000 spectrophotometer. The RNA Integrity Number (RIN) value was calculated using an Agilent Technologies 2100 Bioanalyzer. The mRNA sequencing libraries were prepared according to the manufacturer’s instructions for random hexamer-primed RNA-seq libraries, and were amplified with indexing primers (Illumina Truseq stranded mRNA library prep kit). The quality of the amplified libraries was verified by capillary electrophoresis (Bioanalyzer, Agilent). The RNA-seq libraries were quantified using qPCR with SYBR Green PCR Master Mix (Applied Biosystems). After quantification the libraries were pooled in equimolar amounts and sequenced on an Illumina NovaSeq 6000 system following the provided protocols for 2x100 bp sequencing.

#### *H. rjukjuana* and *P. excavatus* Long-read Sequencing

Using the Covaris G-tube we generated 20Kb fragments by shearing genomic DNA according to the manufacturer’s recommended protocol. Using the AMpureXP bead purification system to remove the small fragments. A total of 5μg for each sample was used as input into library preparation. The SMRTbell library was constructed by using SMRTbell™ Template Prep Kit 1.0 (PN 100-259-100). Using the BluePippin Size selection system we remove the small fragments to create a large-insert library. After sequencing primers were annealed to the SMRTbell template, DNA polymerase was bound to the complex (Sequel Binding Kit 2.0) and purified with SMRTbell Clean Up Columns v2 Kit-Mag: PN 01-303-600. The purification step was performed after polymerase binding to remove excess unbound polymerases and polymerase molecules bound to small DNA inserts. The MagBead Kit was used to bind the library complex with MagBeads before sequencing. The SMRTbell library was sequenced using one Pacific Biosciences 8M SMRT cell usingSequel Sequencing Kit 2.1 on 600 minute movie (continuous long read mode) using the Pacific Biosciences Sequel 2 sequencing platform.

#### *H. rjukjuana* and *P. excavatus* Dovetail Hi-C sequencing

Dovetail HiC libraries were prepared as described previously (Lieberman-Aiden et al. 2009), using a protocol first fixing tissue, then isolating nuclei. The digestion enzyme used was DpnII. The re-ligated DNA was then sheared to ∼350 bp mean fragment size with a Covaris shearing machine, and sequencing libraries were generated using NEBNext Ultra enzymes and Illumina-compatible adapters, then enriched then amplified on streptavidin beads. The libraries were sequenced on an Illumina NovaSeq 6000 system.

#### Haemadipsa rjukjuana and Perionyx excavatus Genome Size Estimation

To estimate the genome size, we used the Illumina whole genome sequencing data to count 17-, 19-, and 21-mers with Jellyfish version 2.1.3 (Marçais and Kingsford 2011). We estimated the genome sizes using GenomeScope (Marçais and Kingsford 2011; Vurture et al. 2017) to obtain estimates for genome sizes, heterozygosity and duplication levels.

#### Haemadipsa rjukjuana and Perionyx excavatus Genome Assembly

For both species, the average coverage of continuous long read (CLR) PacBio sequences was about 96-fold. The average subread length was 16.4 Kb and N50 subread length was 23.2 Kb. The genomes were *De novo* assembled using FALCON-Unzip (Chin et al. 2016) with read lengths greater than the subread N50 value, 23.2 kbp. Duplicate haplotigs were removed using Purge Haplotigs (Roach, Schmidt, and Borneman 2018) with default parameters. The resulting assembly was error-corrected using Pilon (Roach, Schmidt, and Borneman 2018; Walker et al. 2014) with primary contigs to improve the quality of genome assembly results. We checked the assessment of the genome assemblies using BUSCO (Benchmarking Universal Single-Copy Orthologs) (Simão et al. 2015).

#### Haemadipsa rjukjuana and Perionyx excavatus Hi-C Scaffolding Analysis

The Hi-C reads were aligned to primary contigs using juicer version 1.5 (Durand et al. 2016). and the alignments were scaffolded using 3D-DNA (Durand et al. 2016; Dudchenko et al. 2017). The initial scaffolding results were manually reviewed for correcting mis-join and unplaced contigs using Juicebox (Dudchenko et al. 2018). Finally, 3D-DNA was used to regenerate chromosome-level genome assemblies.

#### Haemadipsa rjukjuana and Perionyx excavatus Repeat Annotation

A *de novo* repeat library was constructed using RepeatModeler v.1.0.3 (Bao and Eddy 2002), and RECON/RepeatScout v.1.0.5 (A. L. Price, Jones, and Pevzner 2005), with default parameters. Tandem Repeats Finder (Benson 1999) was also used to predict consensus sequences and classification data for each repeat. All repeats collected by RepeatModeler were searched against the UniProt/SwissProt database (The UniProt Consortium et al. 2022); transposons were excluded. Repetitive elements in the genome were identified using RepeatMasker v.4.0.9 with the repeat library from RepeatModeler.

#### Haemadipsa rjukjuana and Perionyx excavatus Gene Prediction and Annotation

Genome prediction was performed using EVidenceModeler (EVM) v.1.1.1 (Haas et al. 2008), which integrates the results of multiple gene predictions. Repeat-masked genomes were used for *ab initio* gene prediction using GeneMark-ES v.4.68 (Lomsadze et al. 2005) and Augustus v.3.4.0 (Lomsadze et al. 2005; Stanke et al. 2006). Then, the hints for protein and *ab initio* predictions were extracted using protein sequences from Actinopterygii, a clade of bony fishes, in the UniProt/SwissProt protein database (The UniProt Consortium et al. 2022) using ProtHint v.2.6.0 (Brůna, Lomsadze, and Borodovsky 2020). The hints were used to perform protein predictions using GeneMark-EP+ v.4.68 (Brůna, Lomsadze, and Borodovsky 2020) and *ab initio* predictions using Augustus. To obtain transcriptome-level evidence, the PASA pipeline v.2.3.3 (Brůna, Lomsadze, and Borodovsky 2020; Haas et al. 2003) with transcriptome assembly data using Trinity v.2.8.5 (Haas et al. 2013) was used. EVM was used to integrate the *ab initio*, transcriptome, and protein prediction results to obtain the final gene prediction with the weights (ABINITIO_PREDICTION=1, PROTEIN=10, TRANSCRIPT=10). Finally, to predict changes in exons by the addition of untranslated regions (UTRs), the PASA pipeline with Iso-Seq data was used again. Genome Annotation Generator v.2.0.1 (Geib et al. 2018) was used for adding start/stop codon data and generating a gff file. The predicted genes were annotated by aligning them to the NCBI non-redundant protein (nr) database (Geib et al. 2018; Marchler-Bauer et al. 2011) using NCBI BLAST v.2.9.0 (Altschul et al. 1990) with a maximum e-value of 1⨉10^−5^.

### Helobdella austinensis and Hirudo verbana methods

#### *H. austinensis* and *H. verbana* DNA extraction and library preparation

Dovetail Genomics isolated HMW DNA from *Hirudo verbena* testisacs, and from a single clutch of intact, unfed *Helobdella austinensis* individuals. The *Helobdella austinensis* DNA was processed into a whole genome shotgun (WGS) Illumina sequencing library, and was sequenced on a 2x150 run on an Illumina machine to a depth of 678 million read pairs. The *Hirudo verbena* DNA was processed into a Pacific Biosciences SMRT library and was sequenced on CLR mode.

#### H. austinensis and H. verbana genome assembly

The *Helobdella austinensis* WGS Illumina reads were quality trimmed with Trimmomatic (Bolger, Lohse, and Usadel 2014) using the ILLUMINACLIP parameters “2:30:10 SLIDINGWINDOW:13:20 LEADING:20 TRAILING:20 MINLEN:23”. The genome size was estimated as above for *Perionyx excavatus* and *Haemadipsa rjukjuana*, except k-mer sizes of 19, 37, 49, 79, and 109 were used to also infer the k-mer suitability for the Meraculous genome assembler. Meraculous v2.2.4 (Chapman et al. 2011) was used to assemble the genome into contigs and scaffolds using default parameters, a k-mer size of 37, and an estimated genome size of 228 Mbp. Haplotigs were removed from the assembly as part of the Meraculous genome assembly process.

The *Hirudo verbena* Pacific Biosciences CLR reads were assembled using the same procedure as described above for *Perionyx excavatus* and *Haemadipsa rjukjuana*.

### *H. austinensis* and *H. verbana* Dovetail Omni-C Library Preparation and Sequencing

Dovetail “Omni-C” DNAse I Hi-C libraries were prepared by Dovetail Genomics, now Cantata Bio, for both *Helobdella austensis* and *Hirudo verbana* using their provided protocol. The protocol includes chromatin fixation, DNAse I digestion and random re-ligation with biotinylated bridge oligos, and streptavidin enrichment of DNA with re-ligation junctions. The sequencing libraries were generated using NEBNext Ultra enzymes and Illumina-compatible adapters. The libraries were sequenced on an Illumina HiSeqX platform to produce an approximately 30x sequence coverage of the estimated genome size.

#### Helobdella austinensis and Hirudo verbana Assembly Scaffolding with HiRise

The input *de novo* Illumina assembly for *H. austinensis* and the *de novo* Pacific Biosciences *H. verbana* assemblies, and their Dovetail Omni-C library reads, were used as input data for the HiRise genome scaffolder with standard parameters (Putnam et al, 2016). Sequences were curated post-scaffolding by Dovetail genomics such that the assemblies were largely composed of chromosome-level scaffolds.

#### *H. verbana and H. austinensis* genome annotation

The chromosome-scale *Hirudo verbana* genome assembly was annotated by Dovetail genomics. The input data for *Hirudo verbana* annotation hints were existing Illumina RNA-seq data from dissected testisacs, nephridia (Northcutt et al. 2018), ganglia, and neuronal cell types (Heath-Heckman et al. 2021).

The chromosome-scale *Helobdella austinensis* genome was annotated using BRAKER2 v2.1.2 (Brůna et al. 2021), using protein sequences from *Hirudo verbana* as splice site hints.

### Annotation of the *Piscicola geometra* genome

The *Piscicola geometra* genome was downloaded from the Darwin Tree of Life Data Portal (https://portal.darwintreeoflife.org; accession GCA_943735955; “All Seq FASTA”). Because an annotation of this genome was not publicly available and no transcriptome data were available for this species at the time this work was performed, we annotated it using publicly available leech transcriptome assemblies from NCBI TSA (GBRF01.1, GGIQ01.1, GIVY01.1, GIVZ01.1, GIWB01.1, GIWE01.1, GIWF01.1, GIWG01.1, GIWH01.1, GIWI01.1) and transcriptomes from representatives of Piscicolidae and Ozobranchidae. For the latter, raw data were downloaded from NCBI SRA (SRR6766626, SRR10997450, SRR6766627, SRR6766628, SRR6766630, SRR6766631, SRR6766632, SRR10997434, and SRR6766629) and assembled with Trinity v2.8.4 (Haas et al. 2013) with the following optional flags: --trimmomatic --quality_trimming_params “ILLUMINACLIP:/kmk/scripts/trimmomatic/adapters.fasta:2:30:10 SLIDINGWINDOW:4:5 LEADING:5 TRAILING:5 MINLEN:25” --normalize_reads. Assembled transcriptomes were translated using TransDecoder (https://github.com/TransDecoder) following the approach used by (Kutschera et al. 2013; Drábková et al. 2022). Repeats in the genome assembly were annotated and softmasked with RepeatMasker and RepeatModeler (Flynn et al. 2020) using rmblast for both programs (https://www.repeatmasker.org). For RepeatModeler, a maximum genome sample size of 1M and the --LTRStruct option were used. For RepeatMasker, the slow and gccalc options were used. Translated transcriptomes were then mapped to the genome assembly with ProtHint v2.6 (Brůna, Lomsadze, and Borodovsky 2020) with an e-value cutoff of 1⨉10^-25^. Using the output of ProtHint as evidence, annotation of protein-coding genes was performed with BRAKER v2.1.6 (Brůna et al. 2021) with the following settings: “--epmode --softmasking --crf.”

#### Orthology assessment, alignment, and matrix construction

Our approach to generate datasets for phylogenomic analysis followed the bioinformatic pipeline of (Krug et al. 2022). We used OrthoFinder v2.4.0 (Brůna et al. 2021; Emms and Kelly 2019) to identify putatively orthologous sequences among taxa, removing sequences of <100 amino acids from fasta files and keeping the longest non-redundant sequence. Fasta files sampled for ≥75% of taxa were aligned with MAFFT v7.310 (Katoh and Standley 2013), putatively mistranslated regions removed with HmmCleaner (Di Franco et al. 2019), and alignments trimmed to remove ambiguously aligned regions with BMGE v1.12.2 (Criscuolo and Gribaldo 2010). Approximate ML trees were constructed for each alignment with FastTree v2 (M. N. Price, Dehal, and Arkin 2010), and PhyloPyPruner v0.9.5 (https://pypi.org/project/phylopypruner) was used to identify strictly orthologous sequences. We also selected the best 250 genes retained by PhyloPyPruner based on seven properties calculated in genesortR (Mongiardino Koch 2021).

#### Phylogenomic analyses

We performed ML analyses on the complete dataset with IQ-Tree v2 (Mongiardino Koch 2021; Minh et al. 2020) using three strategies: 1) the best-fitting model of amino acid substitution for each partition (-m MFP), 2) Lanfear clustering (-m MFP+merge), and 3) the PMSF model (-m LG+C20+F+G) with the contree produced from the Lanfear clustering analysis specified with “-ft.” The 250-gene dataset was analyzed using the PMSF model with the contree produced by the Lanfear clustering analysis of the complete dataset as the guide tree. Topological support was assessed with 1,000 rapid bootstraps for all analyses. Pairwise patristic distances between taxa were calculated with Patristic v1.0. based on results of the analysis of the complete matrix using Lanfear clustering.

#### Pairwise species comparisons

The odp software v0.3.0 was used to perform reciprocal best blast hits between species pairs, to identify whether a given ortholog corresponded to a known bilaterian, cnidarian, and sponge ancestral linkage group (BCnS ALG), to identify significantly large linkage groups between species, and to create ribbon plots of multiple species. The protocols for these steps have been previously described (Schultz et al. 2023).

#### BLAST searches for Hox genes

We used the TBLASTN operation of BLAST+ 2.15.0 (Camacho et al. 2009) to find reciprocal best hits for Hox genes between an initial set of queries (Hox protein sequences of the polychaete *Capitella teleta*) and a subject genome. To identify potential Hox duplicates, we queried the *Capitella* genome with multiple top hits from the initial subject genome. In an effort to recover Hox genes whose homeoboxes contain introns, we used the TBLASTN commands: -max_intron_length 8000 and -evalue 1e-5.

## Data Availability

Genomes and raw reads will be available under NCBI BioProject PRJNA1109803.

## Code Availability

The odp software, and the scripts used to make dotplots, ribbon diagrams, and to infer ancestral linkage groups, is available on github https://github.com/conchoecia/odp.

## AUTHOR CONTRIBUTIONS

DAW, EACH, CJW, and DHK designed the scientific objectives of the study. KK and DTL conducted phylogenetic analyses. EACH and YSY performed *de novo* genome assembly and annotation. KK, DTS, FO, YSY, CJW, EACH and SJC prepared the genomic data. DTS wrote the code for synteny analyses. DTS and FO conducted the synteny analyses. CJW, EACH and DW performed the Hox evolutionary analysis. DAW, CJW and OS wrote the initial manuscript draft. All authors participated in interpreting the results, editing the manuscript, and creating the figures.

## ETHICS DECLARATIONS

All authors declare that they have no competing interests.

## Supplementary Information

**Supplementary Figure 1:**
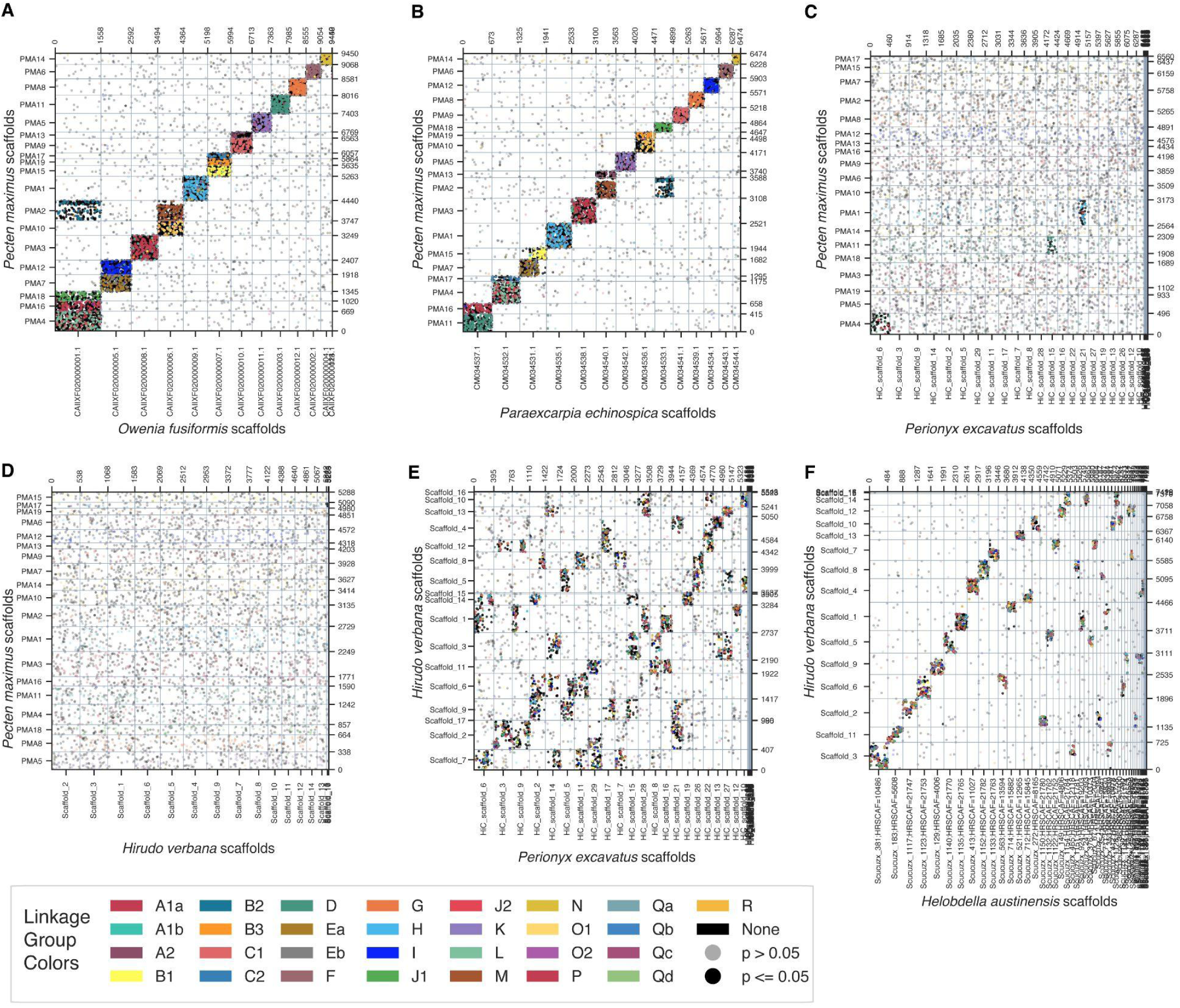
Two-species spiralian protein orthology dotplots. Oxford dotplots showing protein orthology shared between two species, identified with reciprocal best blastp hits by odp. Each axis shows the protein index along each chromosome of the depicted species. The dots are colored if they were identified by odp as being orthologous to a BCnS ALG ortholog. Black dots were orthologs found in these two species, but were not identified, or were a suboptimal hit for, BCnS ALG orthologs. **A.** The scallop *P. maximus* and the annelid *O. fusiformis* have a similar karyotype in which the BCnS ALGs are intact, predominantly on single chromosomes. **B.** The annelid *P. echinospica*, a more recently diverged relative of clitellate annelids, also has a karyotype in which the BCnS ALGs are intact. **C.** The BCnS ALGs have rearranged in the clitellates, shown here in the earthworm *P. excavatus*, and in **D.** the leech *H. verbana*. **E.** Within clitellates the karyotype has changed significantly since the divergence of earthworms and leeches. **F.** Several chromosome fusion and fission events have occurred since the divergence of the leeches *Hirudo* and *Helobdella*.

**Supplementary Figure 2:**
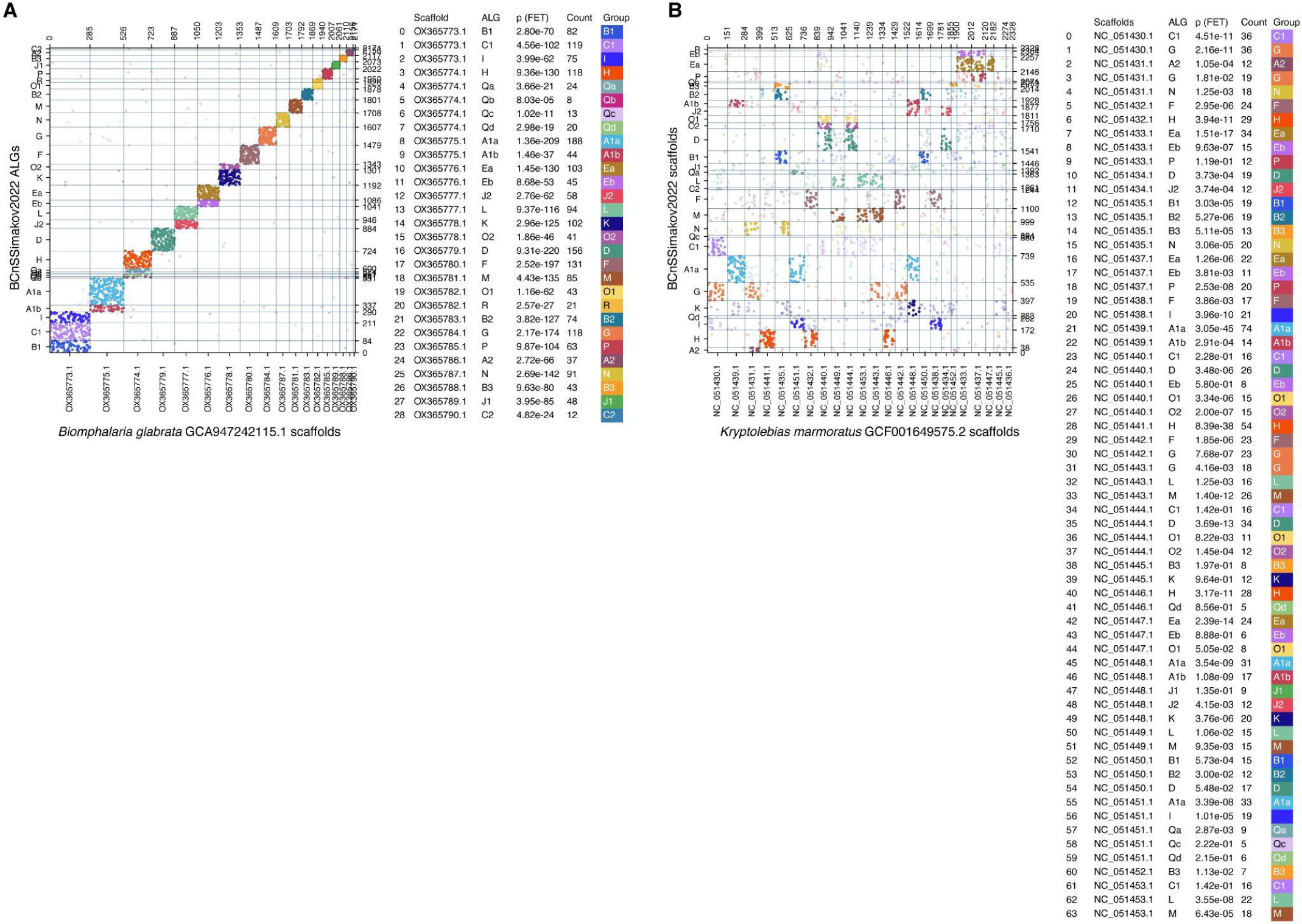
Two-species spiralian protein orthology dotplots. Conservation of ALGs in a simultaneously hermaphroditic mollusk and vertebrate. **A.** The genome of the simultaneously hermaphroditic pulmonate mollusk, *Biomphalaria glabrata*, has a typical spiralian genome (Simakov et al. 2022). **B.** The simultaneously hermaphroditic fish, *Kryptolebias marmoratus*, has a typical vertebrate genome (Simakov et al. 2020).

