## Supplemental Table 1 - docx format for "Acceleration of genome rearrangement in clitellate annelids"

**Supplementary Table 1.** Hox inventories on syntenic genome scaffolds in three clitellate subgroups^1^

| **Comparison of two lumbricid species** | |  |  |  |
| --- | --- | --- | --- | --- |
|  | *Eisenia andrei*^2^ | *Lumbricus rubellus* |  |  |
| Synteny set 1 | Scaffold 1: Post2 | Scaffold 1: [Scr, Antp]^3^, Lox4 Scaffold 14: Lab |  |  |
| Synteny set 2 | Scaffold 2: Pb, Post2, Scr | Scaffold 10: Pb, Post2, Post2 Scaffold 8^4^ |  |  |
| Synteny set 3 | Scaffold 3: Lab, Scr, Lox4, Lox4, Post2, Scr, Lox4, Post2, Lab, Scr, Lox4, Hox3 | Scaffold 3: Post2, Post2, Dfd, Lab, [Lab, Dfd, Lox4, Post2]^7^, Lab, [Scr, Lox2, Hox3] Scaffold 18 |  |  |
| Synteny set 4 | Scaffold 4: Hox3, Scr, Post2, Scr | Scaffold 6: Hox3, Scr, Scr Scaffold 15: Post2 |  |  |
| Synteny set 5 | Scaffold 7: [Lab, Hox3, Scr, Lox5] | Scaffold 2: [Lab, Hox3, Scr, Lox5] |  |  |
| Synteny set 6 | Scaffold 8: Post1 | Scaffold 12: Post1 |  |  |
| Orphans^5^ | Scaffold ctg4208: Scr  Scaffold ctg879: Antp | Scaffold 620: Lox4 |  |  |
| **Comparison of three megascolecid species** | | |  |  |
|  | *Perionyx excavatus*^2^ | *Metaphire vulgaris* | *Amynthas corticis* |  |
| Synteny set 1 | Scaffold 3: Scr, Pb, Post2, Post2 | Scaffold 1: Pb, Lab, Lab, Dfd, Post2, Post2, Lox2, Dfd, Lab, Lab Scaffold 5: Dfd, Dfd, Post2, Post2^6^ | Scaffold 1: Pb, Pb Scaffold 2: Post2, Post2 |  |
| Synteny set 2 | Scaffold 5: Post2 | Scaffold 14: Post2  Scaffold 8 | Scaffold 10 Scaffold 12 |  |
| Synteny set 3 | Scaffold 6: [Lab, Hox3, Scr, Lox5] | Scaffold 2: Lab, Pb, Lox5 Scaffold 3: Lab, Dfd, Lox5 | Scaffold 3: Lox5 Scaffold 6: Lab, Lox5, Dfd, Dfd, Lox5 |  |
| Synteny set 4 | Scaffold 8: [Scr, Antp] | Scaffold 30: [Antp, Scr] Scaffold 37: Scr | Scaffold 18: Antp Scaffold 41 |  |
| Synteny set 5 | Scaffold 10: Post1 | Scaffold 18: Post1 Scaffold 23 | Scaffold 0: Post1 Scaffold 8 |  |
| Synteny set 6 | Scaffold 15: [Post2, Lox4, Dfd, Lab], Lab | Scaffold 1: Pb, Lab, Lab, Dfd, Post2, Post2, Lox2, Dfd, Lab, Lab Scaffold 25: Post2, Lox4, Lab, Lox2, Lox4, Dfd, Hox3, Lab | Scaffold 5: Lab, Dfd, Post2, Post2, Lox4, Lox4, Dfd, Lab Scaffold 35: Lox5, Post2, Lab, Lab, Lox4, Post2 |  |
| Synteny set 7 | Scaffold 17: [Hox3, Lox2, Scr] | Scaffold 6: Scr, Lox2 Scaffold 11: Scr, Hox3 | Scaffold 4: Hox3, Hox3, Scr, Scr Scaffold 13: Hox3, Hox3, Lox4 |  |
| Synteny set 8 | Scaffold 19: Lab | Scaffold 1: Pb, Lab, Lab, Dfd, Post2, Post2, Lox2, Dfd, Lab, Lab Scaffold 41 | Scaffold 5: Lab, Dfd, Post2, Post2, Lox4, Lox4, Dfd, Lab Scaffold 34 |  |
| Synteny set 9 | Scaffold 21: Post2 | Scaffold 28 Scaffold 36 | Scaffold 19 Scaffold 29 |  |
| Synteny set 10 | Scaffold 22: Scr, Hox3 | Scaffold 20: Scr  Scaffold 24: Hox3, Scr | Scaffold 16 Scaffold 25 |  |
| Synteny set 11 | Scaffold 28: Post2, Lox2, Lox4 | Scaffold 31 Scaffold 39 | Scaffold 30 Scaffold 40 |  |
| Synteny set 12 | Scaffold_9 Scaffold_89 | Scaffold 4: Hox3 | Scaffold 19 Scaffold 22 |  |
| Orphans | Scaffold 628: Post2, Lox2, Lox4, Dfd, Lab  Scaffold 629: Lab, Dfd |  | Scaffold 908: Antp |  |
| **Comparison of four leech species** | | | |  |
|  | *Haemadipsa rjukjuana*^2^ | *Helobdella austinensis* | *Piscicola geometra* | *Hirudo verbana* |
| Synteny set 1 | Scaffold 1: [Lab, Scr, Lox5], [Lab, Dfd, Lox4, Post2] | Scaffold 1126^7^: [Lab, Lox5, Scr]  Scaffold 1154: [Lab, Dfd, Lox4, Post2]  Scaffolds 241, 353, 381, 465, 1128 | Scaffold 11: Hox3, [Lab, Dfd, Lox4, Post2]  Scaffold 17: [Lab, Scr, Lox5]  Scaffold 61 | Scaffold 3: [Lab, Scr, Lox5]  Scaffold 14: [Lab, Dfd, Lox4, Post2]  Scaffold 15 |
| Synteny set 2 | Scaffold 3: [Hox3, Lox2, Scr] | Scaffold 712: [Hox3, Scr, Lox2]  Scaffolds 521, 1123, 1130 | Scaffold 16: Scr  Scaffolds 1 and 13 | Scaffold 4: [Hox3, Scr, Lox2]  Scaffolds 6 and 13 |
| Synteny set 3 | Scaffold 9: Scr, Dfd, Lox4, Lox2, Post2, Scr, [Scr, Antp] | Scaffold 149: Scr  Scaffold 183: Scr, [Scr, Antp]  Scaffold 1142: Dfd, Lox4, Lox2, Post2 | Scaffold 5: Dfd, Lox4, Lox2, Post2  Scaffold 8: Antp  Scaffold 15: Scr | Scaffold_9: Dfd, Lox4, Post2  Scaffold_12: Scr  Scaffolds 8 and 11 |
| Orphans |  | Scaffold 1140: Post2 |  |  |

^1^Synteny assignments are based on Oxford dotplots (see Supplementary Figure 1) and genes are listed in their relative order on a scaffold.

^2^Anchor species for the respective subgroup comparison.

^3^Bracketed sets of genes indicate Hox subclusters that are potentially conserved across clitellate subgroups; each of four is marked in a different color.

^4^Gray text indicates a scaffold within a synteny set but with no Hox genes.

^5^Orphan Hox-containing scaffolds could not be assigned to a synteny set due to a paucity of shared orthologs with other leech scaffolds (most are very short Hox-containing scaffolds).

^6^Jin et al. 2020 identified single Post1 and Post2 genes on *Metaphire* scaffold 5 but our analysis found greater support for both being Post2.

^7^This short scaffold is not present in our Oxford dotplots but we include it in leech synteny set 1 given the apparent conservation of a Lab-Scr-Lox5 subcluster in leeches.
